# Validation of DoriVac (DNA origami vaccine) efficacy in a metastatic melanoma model

**DOI:** 10.1101/2024.10.08.617273

**Authors:** Anjali Rajwar, Hawa Dembele, Amanda R. Graveline, Andyna Vernet, Melinda Sanchez, Sarai Bardales, Ick Chan Kwon, Ju Hee Ryu, William M. Shih, Yang Claire Zeng

**Author notes:** These authors contributed equally to this work (Anjali Rajwar, Hawa Dembele).

## Abstract

Metastatic cancer, particularly metastatic melanoma, poses a significant therapeutic challenge due to its resistance to standard treatments and low five-year survival rate of 20–27%. Current therapies show limited success, highlighting the urgent need for novel interventions. Recent advances in cancer vaccine research show great promise in reducing disease recurrence, but these approaches still face challenges pertaining to antigen selection, ease of production, and programmability. To address some of these challenges, we have adapted the DoriVac platform to serve as a cancer vaccine against metastatic melanoma. DoriVac is a DNA origami-based vaccine platform that allows for the codelivery of antigens of choice and the CpG immune adjuvant at an optimal nanospacing to promote Th1 immune polarization. In our study, we observed a significant reduction in lung tumor nodules for mice treated with DoriVac in both the B16OVA and B16F10 melanoma mouse models. DoriVac also appears to be a safe and effective treatment, with no anti-drug antibodies, as indicated by lower anti-dsDNA levels. Importantly, we observed an amplified effect when DoriVac was combined with αPD-L1 immune checkpoint blockade, leading to an even greater reduction in metastatic lung tumor nodules. Our results also indicate an increased activation of antigen presenting cells, NK cells, CD4^+^ T cells and CD8^+^ T cells. These findings suggest that DoriVac, particularly in combination with the αPD-L1 immune checkpoint blockade, can serve as a promising immunotherapy against metastatic cancer.

## Introduction

Malignant melanoma stands out as one of the most aggressive and lethal forms of skin cancer and presents a formidable therapeutic challenge due to its often-limited response to current treatments. According to the American Cancer Society, about 100,000 new melanoma cases are diagnosed each year, with stage IV melanoma carrying only a 20–27% five-year survival rate across cohorts^1^. These discouragingly low survival rates become even more dire considering that recurrence occurs within the first few years of initial treatment due to resistance to conventional therapies such as chemotherapy, radiation, and immune checkpoint blockade inhibitors. These characteristics of metastatic melanoma highlight the urgent need for novel therapeutic approaches ^2,3^.

Cancer vaccines signify a paradigm shift in oncology, harnessing the power of one’s own immune system to recognize and eliminate malignant cells. Cancer vaccines have thus emerged as a groundbreaking strategy for cancer treatment and prevention ^4,5^. Recent advances in this field have shown great success; in particular, Moderna’s personalized mRNA cancer vaccine against high-risk melanoma has shown a significant decrease in disease recurrence in a phase 2 clinical trial when combined with Merck’s Pembrolizumab^6^. These promising results highlight the potential of cancer vaccines as highly effective treatment options against metastatic melanoma. A key consideration in developing therapeutic cancer vaccines is the appropriate delivery of antigens and adjuvants to induce potent T cell responses. One approach to achieve such potent immune responses is the co-delivery of the antigens of interest with an adjuvant of choice. To that end, nanoparticles have emerged as promising cancer vaccine candidates due to their highly programmable nature. An example of such nanoparticles includes DNA origami nanoparticles, which have gained popularity in recent years as a therapeutic platform.

DNA origami utilizes a long single-stranded DNA ‘scaffold’ and numerous short ‘staple’ DNA strands to form functional nanoparticles through Watson-Crick complementary base-pairing^7,8^. DNA nanoparticles can be designed to form pre-programmed shapes of varying sizes and can allow for the arrangement of cargos with nanoscale precision^9,10^. This inherent property enables the manipulation of ligand stoichiometry and facilitates multiplexing^11,12^. In our prior investigation described by Zeng et al., we developed an innovative DNA origami-based cancer vaccine platform (DoriVac) that allows for the codelivery of antigens of choice and the CpG immune adjuvant at an optimal nanospacing to promote Th1 immune polarization^13^. The versatility of this platform was furthermore demonstrated by adapting it as an infectious disease vaccine against diseases such as SARS-CoV-2, HIV, and Ebola^14^. Notably, recent studies from other labs have also demonstrated the utility of DNA origami as a structural framework for the presentation of multiple antigens with controlled stoichiometry and spatial organization, thereby modulating the activation of immune cells ^11,15–19^.

In this study, we aimed to evaluate the efficacy of DoriVac platform against two melanoma tumor models to assess its therapeutic potential in metastatic cancers. We observed significantly reduced tumor burdens in both the B16OVA and B16F10 melanoma models following treatment with DoriVac. This effect was further amplified when DoriVac was combined with αPD-L1 immune checkpoint blockade therapy. Detailed immune cell profiling further disclosed an improved therapeutic effect when combined with αPD-L1 therapy. These findings highlight the potential of the DoriVac platform as an effective treatment for metastatic melanoma and provide a foundation for the development of next-generation cancer immunotherapies.

## Results

### 1. DoriVac treatment mitigates lung metastasis in B16OVA melanoma model

Previous studies demonstrated that the combination of DoriVac with αPD-L1 significantly enhances the therapeutic efficacy of DoriVac in subcutaneous tumor models^13^. This study aimed to evaluate similar effects in a metastatic melanoma model using B16OVA metastasis mouse model. Firstly, we assessed the efficacy of the DoriVac for therapeutic anti-tumor effects in a B16OVA metastasis melanoma mouse model. The metastatic tumor was established by intravenously injecting B16OVA cells.

We fabricated the vaccine using square block DNA origami nanoparticles conjugated with the OVA model antigen and CpG with 3.5 nm spacing as an adjuvant, referred to as the DoriVac **(Figure S1)**. We evaluated various treatment groups: an untreated control, a bolus+SQB group (free SQB nanoparticles, free OVA protein, and CpG), and combinations of αPD-L1 treatment with bolus+SQB or DoriVac. Following tumor inoculation and treatment regimen **(Figure 1A)**, our results indicate a significant decrease in lung tumor nodules for DoriVac-treated groups compared to bolus+SQB or untreated controls **(Figure 1B)**. Additionally, a complementary effect with the αPD-L1 treatment was noted, with the lungs showing significantly reduced tumor burden in these groups **(Figure 1C)**.

**Figure 1:**
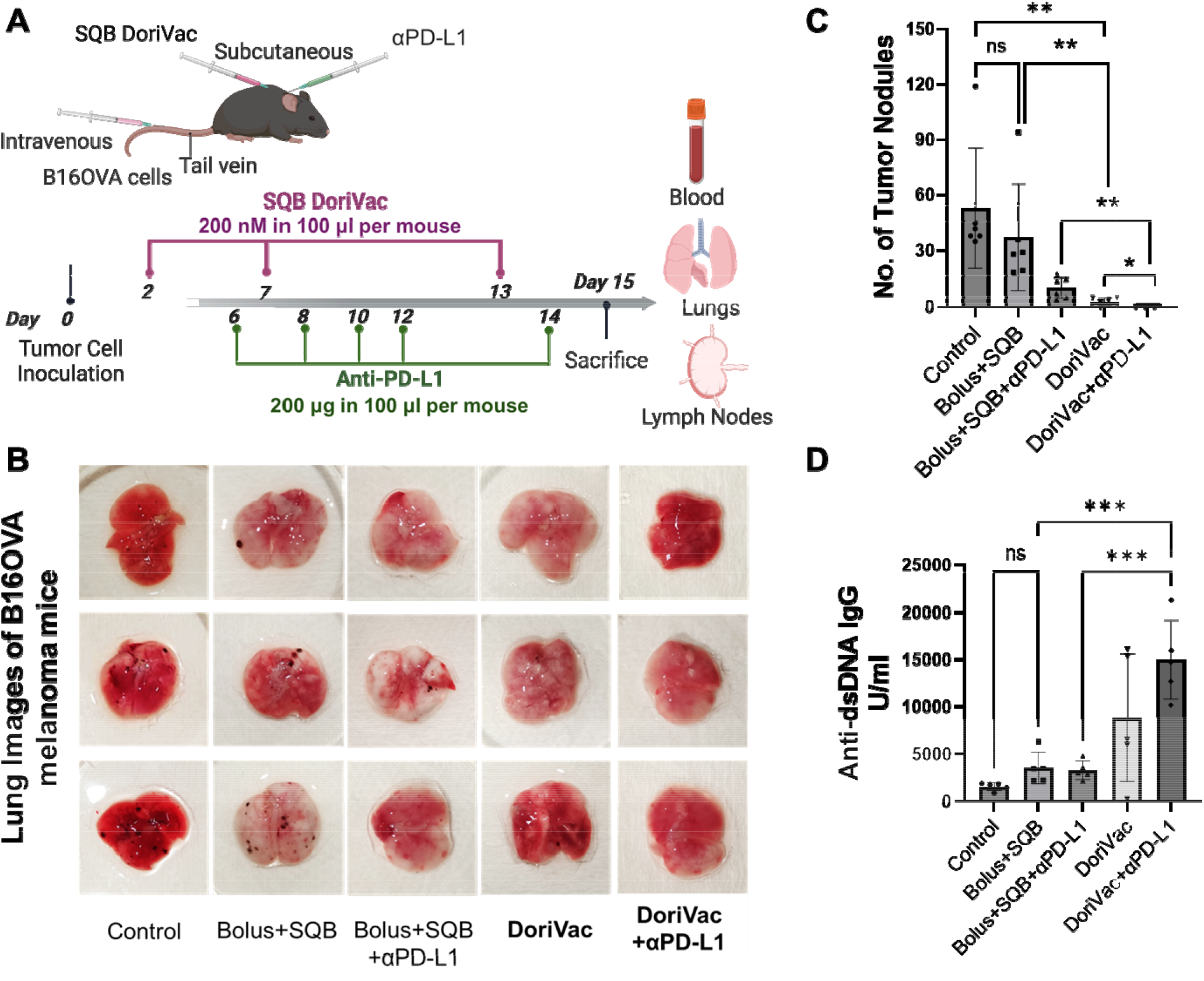
DoriVac treatment reduces tumor burden in the lungs of B16OVA melanoma model. (A) Graphical representation outlining the vaccination regimen employed for the metastasis study, detailing the timing and doses of DoriVac and αPD-L1 treatments. (B) Representative images of the lungs displaying the presence and size of metastatic tumor nodules following the vaccination regimen. (C) Quantification of tumor nodules in the lungs of B16OVA melanoma model across different treatment groups (n=6). (D) Assessment of anti-dsDNA IgG antibodies in the blood samples of different treatment groups (n=5). Error bars indicate the mean with the associated standard deviation (Statistical analysis was performed using one-way ANOVA followed by Tukey’s post hoc multiple comparison test. ***p < 0.001, **p< 0.01, *p < 0.05, ns: not significant, (n = 6).

Using ELISA, we quantified the anti-dsDNA IgG antibody titers in the plasma of various treatment groups to assess the presence of any anti-drug antibodies. The results indicated no significant difference between the control and bolus+SQB groups. However, the DoriVac treatment groups, with and without αPD-L1, exhibited higher titers of anti-dsDNA IgG compared to the untreated control. This suggests that DoriVac, either alone or combined with αPD-L1, elicited a stronger immune response, resulting in increased tumor cell death. Consequently, this increased cell death may have triggered the release of DNA from apoptotic tumor cells, as evidenced by significantly higher antibody titers observed compared to the bolus+SQB group, with or without αPD-L1 (**Figure 1D**). Notably, the bolus+SQB group contained the same amount of SQB nanoparticles, OVA antigen, and CpG as the DoriVac group, suggesting that the elevated anti-dsDNA IgG levels might not be correlated with the DNA origami nanoparticles. Rather, this highlights that the anti-dsDNA IgG antibody response might be associated with tumor-cell killing, with the DNA likely originating from the dead tumor cells.

### 2. DoriVac treatment induces innate immune cell activation

We further investigated the impact of DoriVac on innate immune cells, specifically those that uptake the nanoparticle vaccine and process it for antigen presentation. On day 15 post-inoculation, after euthanizing the mice, we collected the draining lymph nodes for immune cell profiling, where these nanostructures are known to accumulate^13^.

We observed that the treatment with DoriVac led to an increase in the number of activated dendritic cells, shown by higher levels of CD11c^+^CD86^+^ **(Figure 2A, B)**, CD86^+^MHCII^+^ **(Figure 2C, D)**, and CD11c^+^Dec205^+^ **(Figure S2A)** double-positive cells compared to the bolus+SQB vaccine. However, the combination treatment of bolus+SQB vaccine with αPD-L1 did not cause a significant change. Interestingly, when αPD-L1 was given with DoriVac, the effects of extra DC activation induced by DoriVac was attenuated **(Figure 2B, D)**^23,24^. Additionally, our observations revealed that DoriVac treatment resulted in an increase in the population of cells expressing CD40, a co-stimulatory molecule present on DCs known to facilitate the activation of T-helper cells, compared to the bolus+SQB vaccine ^25^ **(Figure 2E)**. However, in combination treatment group of DoriVac with αPD-L1, the previously observed increase in the CD40^+^ cell population was diminished. Notably, a similar reduction was not observed with the combination treatment of bolus+SQB with αPD-L1 **(Figure 2F)**.

**Figure 2:**
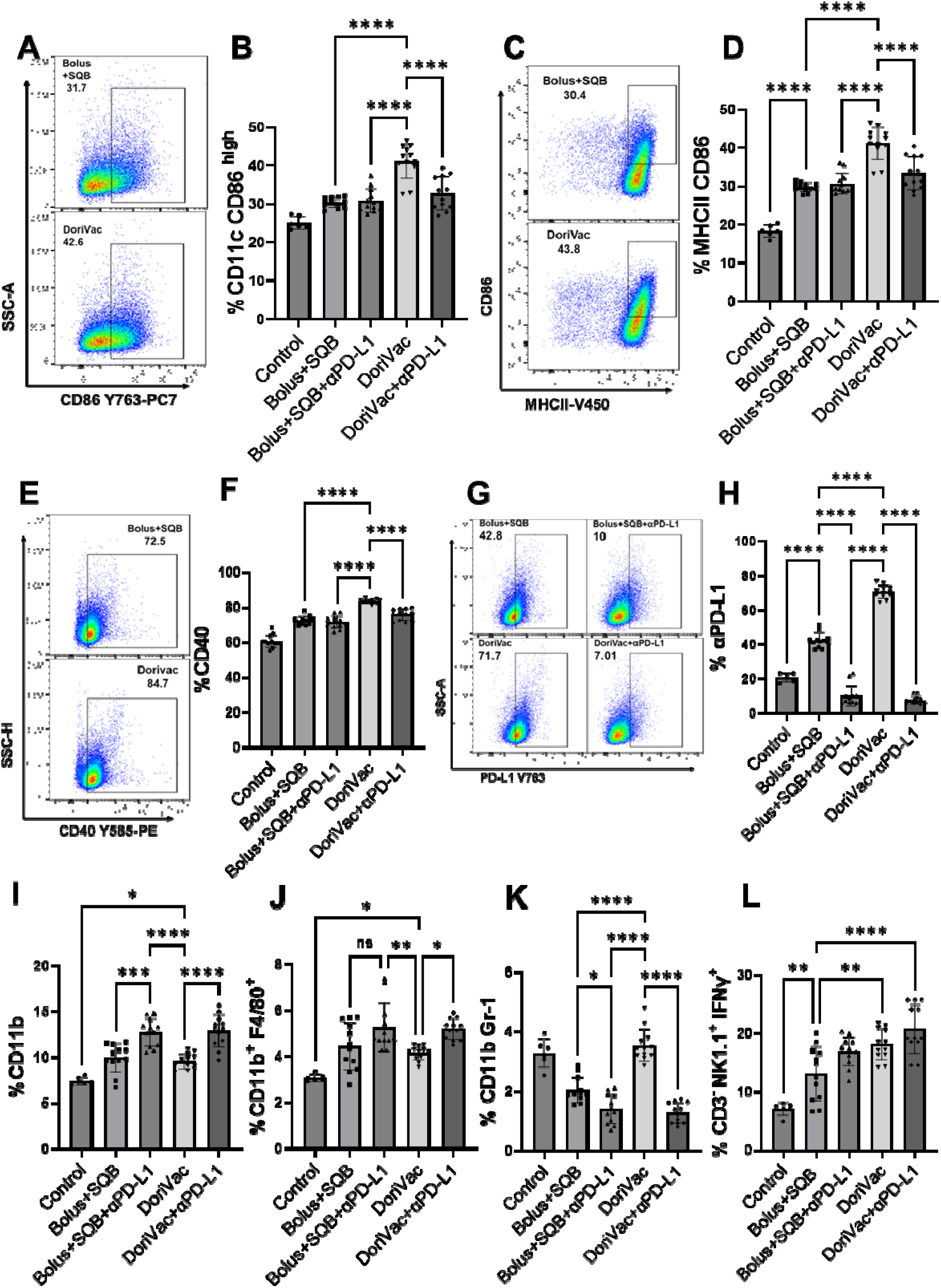
DoriVac stimulates innate immune cell activation in B16OVA melanoma model. (A) Representative flow cytometry scatter plots of Bolus+SQB and DoriVac treatment groups, showing CD11c^+^CD86^+^ dendritic cell population. (B) Quantification of CD11c^+^CD86^+^ dendritic cells as a percentage of total cells in each treatment group. (C) Representative flow scatter plots of Bolus+SQB and DoriVac treatment groups showing the population of activated dendritic cells (MHCII^+^CD86^+^). (D) Quantification graph showing the percentages of MHCII^+^CD86^+^ cells. (E) Representative flow scatter plots of Bolus+SQB and DoriVac treatment groups detecting CD40 co-stimulatory molecules in CD40^+^ cell population. (F) Quantification graph showing the percentages of CD40^+^ cells. (G) Representative flow scatter plots of Bolus+SQB, DoriVac, Bolus+SQB + αPD-L1, and DoriVac + αPD-L1 treatment groups showing αPD-L1^+^ cell population (H) Percentage of αPD-L1+ cells. (I) Percentages of CD11b^+^ monocytes. (J) Percentage of CD11b^+^F4/80^+^ macrophages. (K) Percentage of CD11^+^Gr-1^+^ immunosuppressive cells in mouse lymph nodes as detected by flow cytometry. (L) Percentage of activated natural killer cell subset, identified by the CD3^-^NK1.1^+^IFN γ ^+^ markers. Data were collected from six mice per group. The control groups received no treatment. Error bars indicate the mean with the associated standard deviation. Statistical analysis was performed using one-way ANOVA followed by Tukey’s post hoc multiple comparison test. ****p < 0.0001, ***p < 0.0005, **p< 0.005, *p < 0.01, ns: not significant. Samples from each mouse were analyzed in duplicate for flow cytometry, except for the control group. Therefore, the control group has n=6, while the other groups each have n=12.

Previous studies have elucidated the critical role of PD-L1 expression on DCs in immune checkpoint blockade therapy^26^. DCs serve as the primary antigen-presenting cells for tumor antigen presentation to CD4 T cells and cross-presentation to CD8 T cells. Subsequent upregulation of PD-L1 shields them from cytotoxic T lymphocyte-mediated killing, thereby dampening antitumor responses^27^. However, the majority of previous studies indicated that the blockade of PD-L1 in established tumors promotes the reactivation of tumor-infiltrating T cells, leading to better tumor control^28^. Our DoriVac treatment enhanced PD-L1 expression on DCs,^13^ whereas the combination treatment with αPD-L1 antibody blocked the PD-L1 expression, as depicted in **Figure 2G, H**. Furthermore, the combination therapy of bolus+SQB and DoriVac with αPD-L1 amplified the monocyte as evidenced by CD11b^+^ and mature macrophages by CD11b^+^F4/80^+^ population^29,30^ **(Figure 2I, J)**. We also observed an elevation in the CD11b^+^Gr1^+^ population following DoriVac treatment, suggesting a potential expansion of myeloid-derived suppressor cells (MDSCs), known for their immunosuppressive functions, which could compromise vaccine efficacy^31^. However, in the combination treatment group with DoriVac and αPD-L1, there was a significant decrease in the CD11b^+^Gr1^+^ population **(Figure 2K)**, indicating a potential reversal of MDSC-mediated immunosuppression. The NK-cell activation was characterized by the expression of IFNγ and T-bet in the CD3^-^NK1.1^+^ population **(Figure 2I, and Figure S2C)**. We observed a notable increase in NK cell activation following bolus+SQB treatment compared to the untreated control, with an even more pronounced increase in the DoriVac treatment group. Moreover, the combination treatment of bolus+SQB and DoriVac with αPD-L1 further increased this activated NK cell population.

### 3. Activation of cell-mediated immunity by DoriVac and αPD-L1 combination treatment

We next conducted lymphocyte immune profiling to evaluate the impact of DoriVac combined with αPD-L1 therapy on cell-mediated immunity. In the combination treatment group of bolus+SQB and DoriVac with αPD-L1, we observed a marked increase in the populations of PD1^+^CD4^+^ **(Figure 3A, B)** and CD4^+^IFN γ ^+^ **(Figure 3C)** cells compared to the groups treated with bolus+SQB or DoriVacalone. This suggests enhanced immune activation due to complementary effects of the combined therapy. The blockade of the PD-1/PD-L1 pathway with αPD-L1 likely contributed to the activation of CD4^+^T cells and subsequent increase in IFNγ production, highlighting the potential of this combined treatment in promoting anti-tumor immune responses.

**Figure 3.**
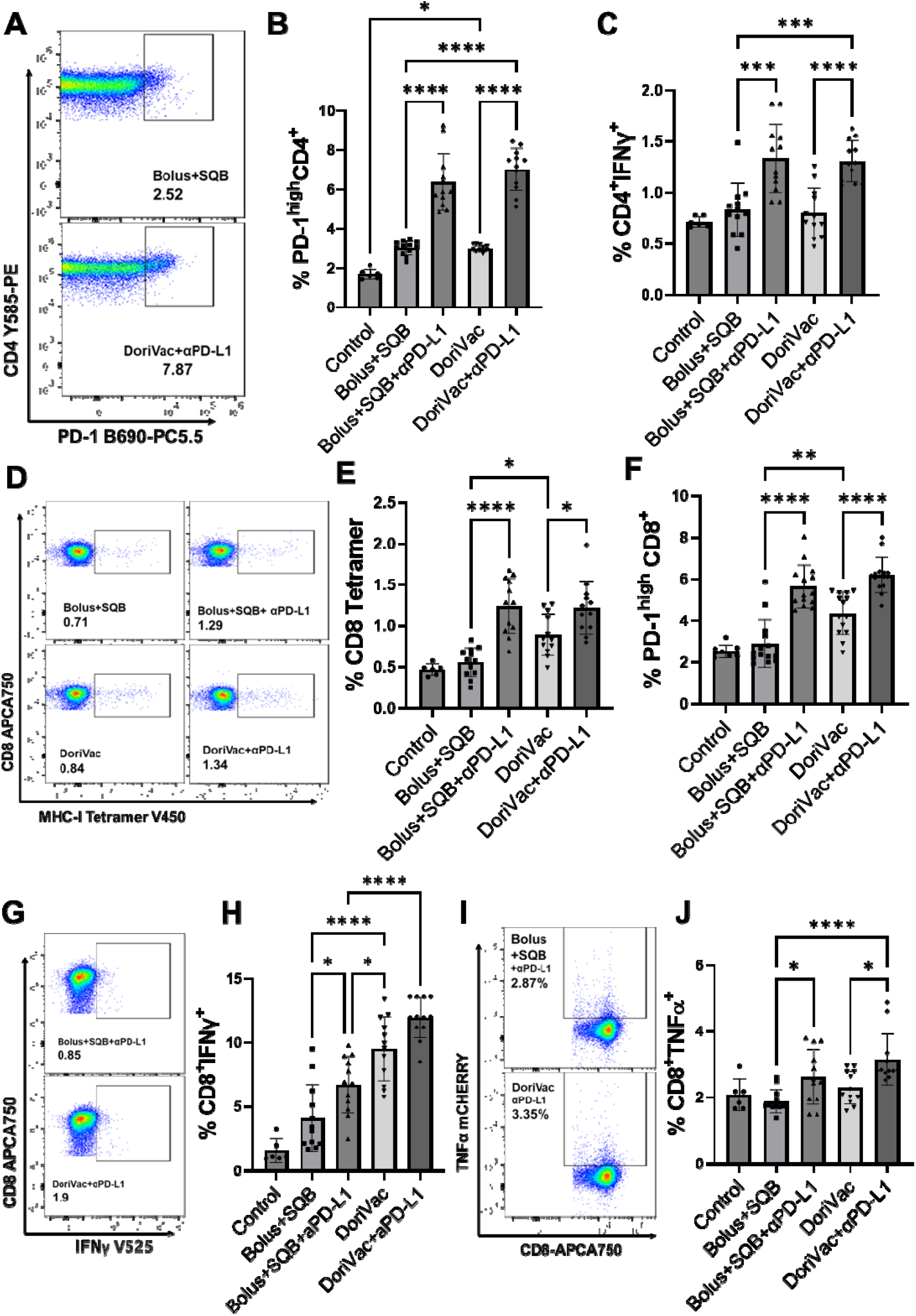
αPD-L1 enhances cell-mediated immunity induced by DoriVac in B16OVA model. (A) Representative flow scatter plots of Bolus+SQB and DoriVac+αPD-L1 treatment groups, showing CD4 T cells expressing the immune checkpoint protein PD-1. (B) Quantification graph showing the percentages of CD4^+^PD-1^high^ T cells in each group. (C) Percentage of IFNγ-producing CD4^+^ cells in the treatment groups. (D) Representative flow scatter plots showing CD8 T cells bound to tetramer molecules, identifying antigen-specific CD8 T cell population. (E) Percentage of CD8^+^Tetramer^+^ antigen-specific T cells. (F) Percentage of CD8 T cells expressing surface marker PD-1 in the treatment groups. (G) Representative flow scatter plots of Bolus+SQB and DoriVac treatment groups showing CD8^+^IFNγ^+^ T cell population. (H) Percentage of IFNγ-secreting CD8 T cells. (I) Representative flow scatter plots of Bolus+SQB+αPD-L1 and DoriVac+αPD-L1 treatment groups showing CD8^+^TNF α ^+^ T cell population. (J) Percentage of CD8^+^TNFα^+^ T cells. The cells were isolated from mouse lymph nodes and analyzed by flow cytometry. Data were collected from six mice per group. The control groups received no treatment. Error bars indicate mean with the associated standard deviation. Statistical analysis was performed using one-way ANOVA followed by Tukey’s post hoc multiple comparison test. ****p < 0.0001, ***p < 0.0005, **p < 0.001, *p<0.05. Samples from each mouse were analyzed in duplicate for flow cytometry, except for the control group. Therefore, the control group has n=6, while the other groups each have n=12.

Additionally, there was a substantial increase in PD1^+^CD8^+^ **(Figure 3F)** and CD8 tetramer cell populations. The increase in PD1^+^CD8^+^ cells observed in the combination treatment groups of bolus+SQB and DoriVac with αPD-L1, compared to bolus+SQB and DoriVac alone, suggests enhanced T-cell activation and effector function against tumor cells. Similarly, the rise in CD8 tetramer cells in the combination treatment groups with αPD-L1 indicates improved priming and activation of antigen-specific CD8 T cells, facilitating the recognition and elimination of tumor cells. **(Figure 3D, E)**. Furthermore, we noted a significant increase in CD8^+^IFNγ^+^ **(Figure 3G, H)** and CD8^+^TNFα^+^ **(Figure 3I, J)** cell subsets in the combination treatment group of DoriVac with αPD-L1 compared to the bolus+SQB group, indicating heightened cytotoxic activity and increased pro-inflammatory cytokine production by CD8 T cells. These effector functions of CD4 and CD8 T cells are known to be crucial for directly targeting and eliminating tumor cells, as well as for creating an inflammatory microenvironment conducive to anti-tumor immune responses.

### 4. Efficacy of DoriVac treatment in the B16F10 metastatic melanoma model

We next assessed the efficacy of DoriVac in a wild type B16F10 metastasis melanoma model, known for its aggressiveness and its status as a “cold” tumor model with high metastatic potential. Despite its lack of immune cell infiltration, the B16F10 model closely mirrors key aspects of human melanoma, making it a valuable tool for preclinical investigations into melanoma biology. To establish the metastatic spread, we developed a B16F10 metastatic tumor model using a similar approach used for the B16OVA model, inoculating 50,000 cells via the tail vein to mimic metastatic melanoma progression in mice. The vaccine formulation included SQB NP, which were co-cultured with B16F10 tumor supernatant that had undergone immunogenic cell death to capture tumor-specific antigens **(Figure S3)**. In this model, SQB captured a broader spectrum of antigens from the tumor cell supernatant, resulting in a wider antigenic load compared to the B16OVA model, which involves only the OVA antigen.

Our study included four treatment groups: untreated control, αPD-L1 monotherapy, DoriVac monotherapy, and combination therapy with αPD-L1 and DoriVac. The mice received three doses of vaccine on days 2, 7, and 13, followed by five doses of αPD-L1 after vaccination on days 3, 8, 10, 12, and 14. Subsequently, the mice were euthanized on day 15, and blood, lungs, and lymph nodes were harvested for analysis **(Figure 4A)**. We observed metastasis in the lungs, as indicated by the presence of black colored tumor nodules. Lung samples appeared cleaner and exhibited significantly fewer tumor nodules in the DoriVac and combination treatment groups compared to both the untreated control and αPD-L1 monotherapy groups **(Figure 4B)**. DoriVac treatment, whether administered alone or in combination with αPD-L1, resulted in a significant reduction in tumor nodules compared to αPD-L1 monotherapy **(Figure 4C)**. Immune cell profiling revealed increased populations of innate immune cells, marked by CD11c^+^ dendritic cells, in the combination treatment group **(Figure 4D, E)**. However, we observed an increase in activated CD11c^+^CD86^+^ dendritic cells upon DoriVac treatment, which was attenuated by the combination treatment with αPD-L1 **(Figure 4F, G)**. Similar to the B16OVA model, we observed an increase in the CD11c^+^CD40^+^ and CD11b^+^MHCII^+^CD86^+^ cell populations upon DoriVac treatment. However, this effect was attenuated when combined with αPD-L1 treatment, while no significant changes were observed with αPD-L1 monotherapy **(Figure S4A, S4B)**. Additionally, we observed an increase in CD11b^+^ population in the DoriVac group **(Figure 4H, I)**. We also observed a significant increase in macrophage markers F4/80 and CD64 in the DoriVac group **(Figure 4J, K, and Figure S4C)**. However, this effect was reversed by the combination treatment with αPD-L1 in B16F10 model. The differences in C11b and F4/80 markers between the B16F10 and B16OVA models might be attributed to the timing of αPD-L1 administration. The early intervention of αPD-L1 on Day 3 in the B16F10 model likely influenced the initial immune response, leading to a distinct pattern of immune cell activation and differentiation compared to the later administration on Day 6 in the B16OVA model.

**Figure 4:**
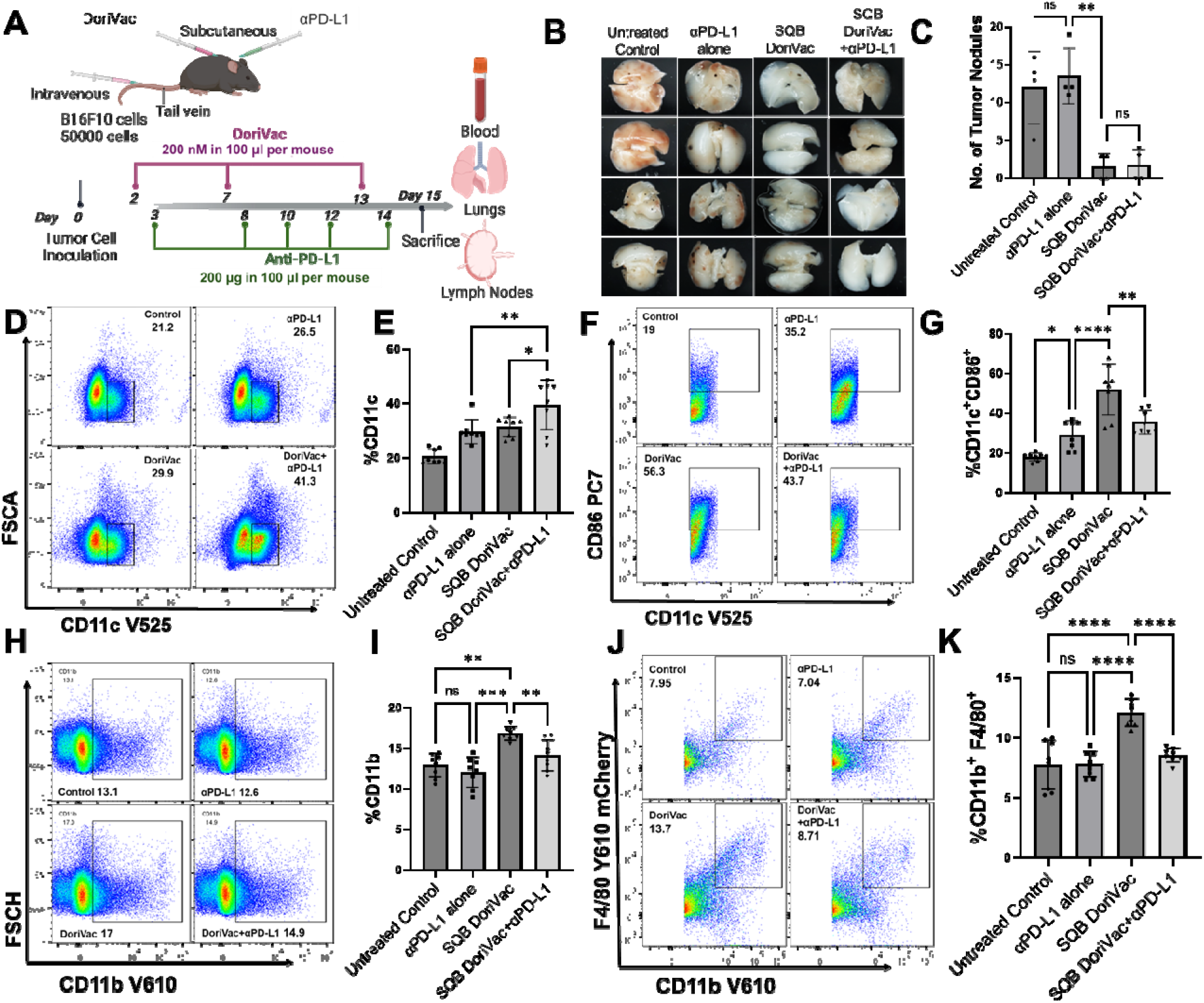
DoriVac treatment reduces metastasis and activates innate immune cells in the B16F10 metastasis model. (A) Schematic representation outlining the vaccination regimen employed for the B16F10 metastasis study, detailing the timing and doses of DoriVac and αPD-L1 treatments. (B) Representative images of the lungs displaying metastatic tumor nodules from different treatment groups. (C) Quantification of metastatic tumor nodules in the lungs from each treatment group. Representative flow scatter plots of different treatment groups showing (E) CD11c^+^ dendritic cell population, (F) CD11c^+^CD86^+^ activated dendritic cells, (H) CD11b^+^ monocyte population, and (J) CD11b^+^F4/80^+^ macrophages isolated from lymph nodes. Quantification graph showing the percentages of (E) CD11c^+^ dendritic cells, (G) CD11c^+^CD86^+^ activated dendritic cells, (I) CD11b^+^ monocytes, and (K) CD11b^+^F4/80^+^macrophages, respectively. The cells were isolated from mouse lymph nodes and were analyzed by flow cytometry. Data were collected from four mice per group. The control groups received no treatment. Error bars indicate the mean with the associated standard deviation. Statistical analysis was performed using one-way ANOVA followed by Tukey’s post hoc multiple comparison test (****p < 0.0001, ***p < 0.001, **p < 0.01, *p<0.05). Samples from each mouse were analyzed in duplicate for flow cytometry, therefore each group had n=8.

### 5. DoriVac treatment modulates CD4 and CD8 T cell populations

In the B16F10 melanoma model, we observed the significant increase in CD4^+^PD-1^+^ population upon DoriVac treatment **(Figure 5A**. This increase was further amplified in the αPD-L1 combination group. This finding suggested a complementary interaction between DoriVac and αPD-L1 therapies, enhancing CD4 T cell activation beyond what was observed with DoriVac monotherapy. Additionally, we observed an increase in regulatory T cells (Tregs), as evidenced by increases in CD4^+^CD25^+^FoxP3^+^ with DoriVac treatment **(Figure 5B)**. Tregs are a specialized subset of CD4^+^ T cells that play a crucial role in maintaining immune tolerance and preventing autoimmune responses by suppressing the activation and proliferation of other immune cells^32^. Conversely, monotherapy with αPD-L1 did not exhibit significant impact on the Treg population, suggesting the moderate immune cell activation. However, a combination therapy comprising DoriVac with αPD-L1 elicited a further enhancement in the Tregs **(Figure 5C)**, indicating an enhanced immune cell activation. DoriVac treatment also resulted in an increase in the CD4 T cell expressing interferon-gamma (IFNγ) and CD107a, a marker indicating degranulation and cytotoxicity in T cells^33^ **(Figure 5D, E**,**F)**. The combination of DoriVac with αPD-L1 treatment further intensified the responses, suggesting a complementary effect between the two treatments in enhancing immune activation

**Figure 5:**
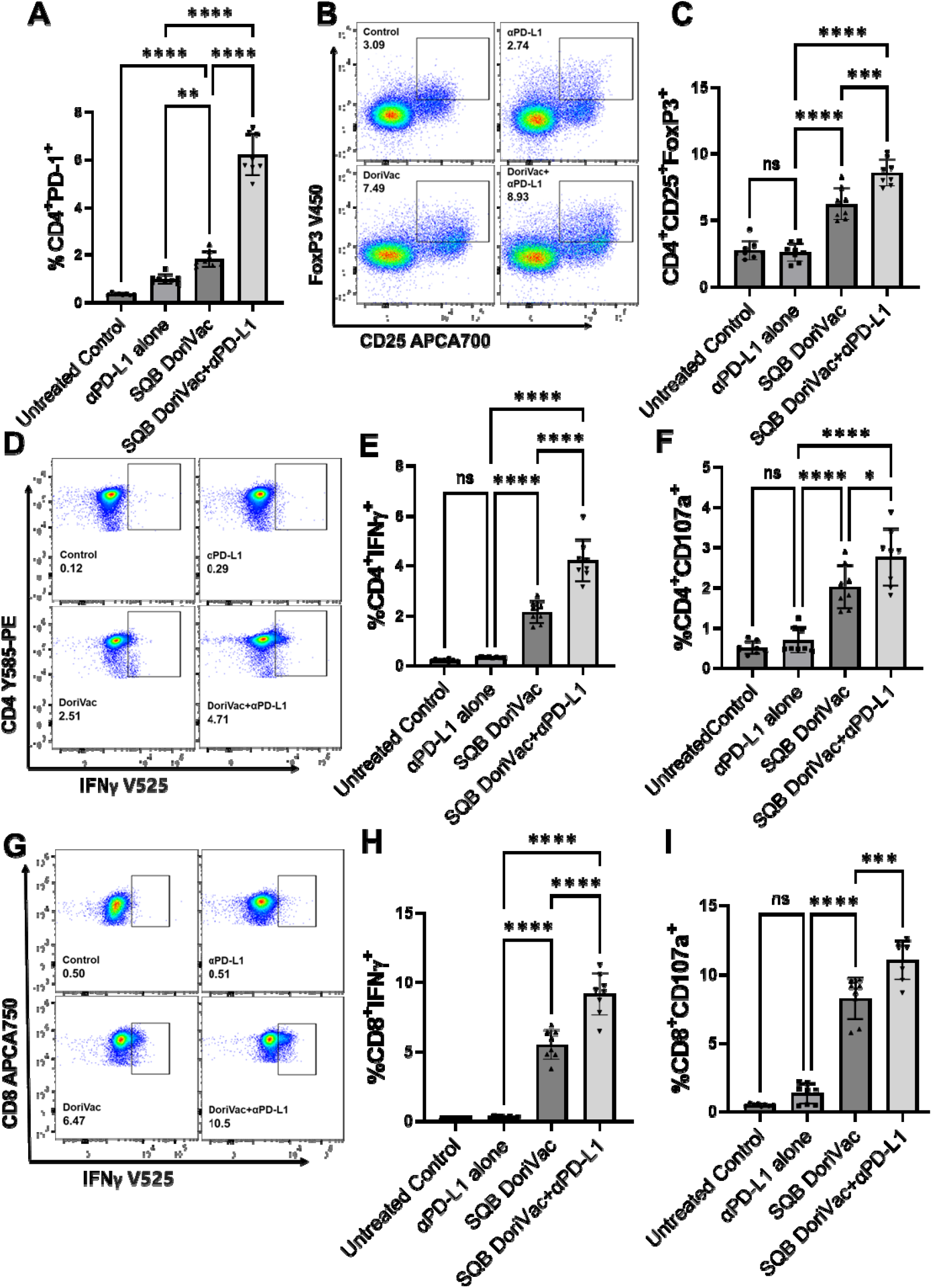
DoriVac and αPD-L1 enhance the activation of the CD4 and CD8 T cell populations. (A) Percentages of CD4^+^ PD-1^+^ T cells in the different treatment groups, showing the activation status of CD4^+^ T cells. (B, C) Representative flow cytometry scatter plots and corresponding quantification of regulatory T cells (Tregs) from different treatment groups, highlighting CD4 T cells co-expressing CD25 and FoxP3. (D, E) Representative flow cytometry scatter plots and quantification of CD4^+^IFN-γ ^+^ T cells, showing IFN-γ producing CD4^+^ T cells. (F) Percentages of CD4^+^CD107a^+^ T cells, indicating degranulation and cytotoxic activity of CD4^+^ T cells. (G, H) Representative flow cytometry scatter plots depict CD8^+^IFN-γ ^+^ T cells in mouse lymph nodes demonstrating the activation of cytotoxic CD8^+^ T cells, along with corresponding quantification. (I) Percentages of CD8 T cells expressing the CD107a marker of degranulation, indicative of cytotoxic T cell activity. Data were collected from four mice per group. Untreated control groups received no treatment. Error bars represent the mean ± standard deviation. Statistical analysis was performed using one-way ANOVA followed by Tukey’s post hoc multiple comparison test. (****p < 0.0001, ***p < 0.001, **p < 0.01, *p<0.05, ns: not significant. Samples from each mouse were analyzed in duplicate for flow cytometry, therefore each group had n=8.

Similar to CD4^+^ T cell activation, we also observed the increase in CD8^+^CD107a^+^ cell population post DoriVac treatment, suggesting heightened cytotoxic activity of CD8 T cells, which was further amplified with the combined treatment of DoriVac with αPD-L1 **(Figure 5G, H)**. Similar trends were observed with the CD8^+^IFNγ^+^ population, demonstrating an increase following DoriVac treatment alone and a further augmentation in combination treatment of DoriVac with αPD-L1, but not with αPD-L1 monotherapy **(Figure 5I)**.

Moreover, we also observed that DoriVac treatment showed a substantial increase in the memory helper T cells, identified by the expression of CD4^+^CD44^+^ **(Figure S5A)** and memory cytotoxic T cell subsets CD8^+^CD44^+^ **(Figure S5B)**, compared to αPD-L1 monotherapy, with further enhancement seen in the combination therapy of DoriVac with αPD-L1. This suggests DoriVac’s antigenic stimulation likely promoted CD4 and CD8 T cell activation and proliferation with a memory phenotype (CD44^+^). Conversely, the population of CD4 and CD8 T cells expressing tumor necrosis factor-alpha (TNF-α) exhibited an increase following combination treatment of DoriVac with αPD-L1, indicating a distinct inflammatory response potentially beneficial for antitumor immunity. Intriguingly, no significant change was noted in this population with DoriVac monotherapy, suggesting a unique influence of DoriVac combination therapy with αPD-L1 in eliciting TNF-α expression **(Figure S5C, 5D)**.

Overall, these results suggest that DoriVac enhances cytotoxic CD4 and CD8 T cell activity more effectively than αPD-L1 monotherapy, with further amplification observed when DoriVac was combined with αPD-L1.

## Methods

### 1. Vaccine Fabrication

The SQB DNA origami nanoparticle used as a vaccine in study was repurposed from a study previously published by our lab^20^. The p8634 scaffold strand used for the SQB was produced in-house as previously described^21^, and the staple strands were purchased from IDT. The SQB origami was folded using an 18-hour annealing program as previously described in a buffer containing 5mM Tris HCl, 1 mM ethylenediaminetetraacetic acid (EDTA; pH 8.0), and 12 mM MgCl_2_. The reaction conditions contained a scaffold concentration ranging from 50 to 100 nM, with 5-to 20-fold molar excess of the staple strands. The fully folded origami was purified using PEG precipitation and analyzed using 2% agarose gel electrophoresis. The structural integrity of the SQB origami was confirmed via Transmission Electron Microscopy (TEM) using 0.75% (w/v H_2_O) uranyl formate and negative staining techniques. To ensure the stability of the SQBs in physiological conditions, oligolysine-PEG5k (K10-PEG5k) coating was employed. Briefly, an appropriate amount of K10-PEG5k to neutralize the charge on the surface of the SQB was added and the mixture was placed on the shaker at 37°C for 30 min. The final product was stored at -20°C until injection.

### 2. Antigen preparation

#### *B16OVA* melanoma model vaccine

The OVA protein was first attached to an oligonucleotide using a succinimidyl 4-(N-maleimidomethyl) cyclohexane-1-carboxylate (SMCC) linker and purified using a NAP column (GE Healthcare Life Sciences). Unconjugated oligonucleotides were removed by 30K Amicon filtration (Sigma). The resulting OVA-oligo was conjugated onto the SQB origami surface using DNA handle/anti-handle hybridization. Conjugated OVA-oligos were incubated with the SQB origami nanoparticle at 37°C for 2 hours in three-fold molar excess, followed by removal of unbound OVA-oligo using PEG precipitation. The resulting OVA-oligo was then incubated with the purified SQB origami at 37°C for 2 hours in a three-fold molar excess of OVA-oligo. Unbound OVA-oligos were removed using PEG precipitation.

#### *B16F10* melanoma model antigen capture vaccine

##### SQB fabrication for antigen capture

The SQB origami nanoparticles were fabricated and purified as described above. A hydrophobic coiled-coil peptide^22^ was conjugated onto the SQB origami surface using DNA handle/anti-handle hybridization. The coiled-coil peptide was first attached to an oligonucleotide using DBCO-Azide copper free click chemistry: The amine-modified oligonucleotide was reacted with Dibenzocyclooctyne-N-hydroxysuccinimidyl ester (DBCO-NHS ester; Millipore, #761524) in phosphate buffer pH 8.0 overnight, purified via NAP Column (GE Healthcare Life Sciences, #17-0852-02) and ethanol precipitation. Briefly, the NAP purified oligo-DBCO was mixed with 100 proof ethanol to achieve an 80% ethanol concentration and Sodium Acetate in a 1:10 v:v ratio. The resulting mixture was placed at -80°C for 1 hour and spun at 16000g for another hour. After spinning, the supernatant was discarded, and the pellet obtained was washed twice using 75% ethanol. The oligo-DBCO obtained was then mixed with azide-modified peptide in a 2:1 peptide:oligo ratio and incubated overnight at room temperature. The peptide-oligonucleotide was purified using 8% PAGE gel extraction, filtering through Freeze ‘N Squeeze DNA Gel Extraction Spin Columns (Bio-Rad; #7326165) and then using ethanol precipitation. The purity of the sample was confirmed by running a 15% dPAGE gel with the unpurified and purified sample. Peptide-oligos were then incubated with purified SQB origami particles at 37°C for 1–2 hrs and the excess peptide-oligo was removed via PEG precipitation.

##### Tumor cell irradiation and co-culture with SQB

B16F10 melanoma cells were cultured at 37°C in DMEM media supplemented with 10% heat inactivated FBS and 1% penicillin-streptomycin. Once the cells reached full confluency, the cells were washed one with PBS and then resuspended in PBS. The B16F10 cells were then irradiated using a gamma irradiator for 72.5 minutes to reach 100 Gy photons. The cells were spun down, then resuspended in serum-free RPMI media and incubated at 37°C for 48 hours. Following 48 hours of incubation, the resulting culture was spun down, and the protein-rich cell supernatant was collected and concentrated using a 3k MWCO Amicon filter.

### 3. Silver staining

Samples were run on a SDS PAGE gel and stained using silver staining following DNase degradation to confirm antigen conjugation. The DNase digestion was performed as follows: 1 µg of DNA origami nanoparticle was mixed with 1 U/µl DNase I (2,000 units/mL, New England Biolabs, M0303S) mixed with 10× DNase I buffer diluted in water (Gibco). The resulting solution was then incubated for 30 min at 37°C. Following digestion, the sample was mixed with 4× LDS NuPAGE loading buffer (ThermoFisher; #NP0008) and incubated at 95°C for 2 min. The mixture obtained was loaded on a 4– 12% NuPAGE Bis-Tris gels (ThermoFisher; #NP0322) and ran in 1× MES SDS buffer (ThermoFisher; #NP0002) at 150V for 45 minutes. The gel was then subjected to silver staining as described by the manufacturer (Pierce, #24612). The gel obtained was imaged using the Silver Stain setting on Image Gel 6 on Gel Doc EZ Imager (Bio-Rad).

### 4. Animal model

The B16OVA and B16F10 melanoma cells used in this study were a kind gift from Dr. David Mooney (Harvard University). Both cell lines were cultured at 37 °C in DMEM media supplemented with 10% heat inactivated FBS, as well as 1% penicillin and streptomycin. Prior to tumor inoculation, the appropriate number of cells for each model was trypsinized and resuspended in PBS. The cell suspensions were then injected through the tail vein following an IACUC approved protocol. All mice were euthanized on day 15 after initial tumor inoculation for subsequent analysis.

### 5. Vaccination regime

#### *B16OVA* melanoma model

Six to eight weeks old female C57BL/6 mice were injected through the tail vein with 300,000 B16OVA cells in 200 μL suspension. The mice were then vaccinated at days 2, 7, and 13 following initial tumor inoculation. The mice in the combination treatment groups received αPD-L1 antibodies on days 6, 8, 10, 12 and 14 following initial tumor inoculation. On day 15, all animals were sacrificed and their heart blood, lymph nodes, and lungs were harvested for further processing.

#### *B16F10* melanoma model

Six to eight weeks old female C57BL/6 mice were injected through the tail vein with 50,000 B16F10 cells. The mice were then vaccinated at days 2, 7, and 13 following initial tumor inoculation. The mice in the combination treatment groups received αPD-L1 antibodies on days 3, 8, 10, 12, and 14 following initial tumor inoculation. On day 15, all animals were sacrificed and their heart blood, lymph nodes, and lungs were harvested for further processing.

### 6. Lung isolation and tumor nodules quantification

Following euthanasia, the lungs were perfused with 20 mL of PBS through the heart to clear out most of the blood. The lungs were then harvested and immediately immersed in a 35 mm petri dish containing cold PBS. For long term storage, lungs were moved to vials containing formalin. For lung imaging, the lungs were moved to a dry 35 mm petri dish and the tumor nodules on individual lungs were counted manually.

### 7. Processing Lymph Nodes

The left axillary and superficial cervical lymph nodes of the euthanized mice were harvested and stored in cold PBS as previously described (insert citation). The lymph nodes were then processed into single cell suspensions through physical dissociation using the back of a syringe plunger. The resulting mixture was then passed through a 70 µm strainer and washed with PBS prior to plating for flow processing.

### 8. Flow cytometry

After plating of the lymph node single cell suspensions, the cells were stained with Zombie UV viability dye (BioLegend; #423108) and washed with cell staining buffer (BioLegend, #420201). Cell surface antibodies were then added to the cells following the manufacturer’s protocol. For intracellular staining, permeabilization and fixation reagents (BioLegend; #424401) were used prior to adding the antibodies. The stained cells were then fixed and analyzed on a Cytoflex LX flow cytometer. The resulting data was then analyzed using FlowJo (https://www.flowjo.com/). For flow analysis, the gating was performed excluding debris, doublets, and dead cells. Single and double positive populations were gated out of the live cell population first, then out of other mother populations.

### 9. Toxicity

Following mice euthanasia, heart blood was collected in heparin-rinsed tubes. The blood-filled tubes were spun down at 4C for 5 min to separate the plasma from cells. The collected plasma was then incubated with anti-mouse dsDNA IgG (#5120), anti-mouse PEG IgG (#PEG-030), and anti-mouse dsDNA IgA/G/ (#5110) antibodies. For anti-dsDNA antibody tests, the plasma samples were diluted 100 times. For anti-PEG antibody tests, the plasma samples were diluted 10 times. The ELISA experiment was performed as described by the manufacturer (Alpha Diagnostic International).

### 10. Statistical analysis

GraphPad Prism 10 was utilized for graph creation and p-value calculation. Experimental procedures were not randomized, and investigators were not blinded to allocation during both experimentation and outcome evaluation. Mean values along with their associated standard deviations (SD) were employed and mentioned accordingly. The statistical significance of all flow and ELISA data in Figures 2–6 was determined using unpaired t-tests. A p-value of ≤ 0.05 was considered statistically significant. Error bars represent standard deviation (SD).

## Conclusion

In this study, we evaluated the efficacy of the DoriVac cancer vaccine platform against two metastatic melanoma models, B16OVA and B16F10. Our results indicate that DoriVac effectively mitigated metastasis while generating robust innate and cell-mediated immune responses. Specifically, our results indicate that DoriVac treatment significantly activated innate immunity through an increase in the activation of CD11c^+^ dendritic cells, as well as other CD11b^+^ myeloid cell populations such as monocytes and macrophages in both the B16OVA and B16F10 models. Bridging the innate and adaptive immunity, DoriVac significantly induced cellular immunity as indicated by the activation of CD4^+^ and CD8^+^ T cells through increased secretion of TNFα and IFNγ, highlighting the induction of a strong anti-tumor response in both models.

In the B16OVA model, which features a defined antigen allowing for the detection of tetramer-expressing CD8 T cell population, DoriVac-treated groups showed a notable increase, suggesting an antigen-specific immune response. We also observed NK cell activation, consistent with previous findings on for DoriVac^13^. We investigated the effects of αPD-L1 on the immune cell activation, both as monotherapy and in combination with DoriVac. Our findings indicated that DoriVac alone was more successful in eliciting a potent immune response compared to bolus+SQB vaccine or αPD-L1 monotherapy. Furthermore, the combination of αPD-L1 with DoriVac enhanced the NK and T cell immune responses beyond what was observed with either monotherapy. Importantly, for the treatment condition of empty DoriVac (i.e. bolus+SQB), we did not observe the production of anti-drug antibodies, as evidenced by the anti-dsDNA levels, suggesting that DoriVac can serve as a safe and effective treatment option. Collectively, these findings highlight the efficacy of the DoriVac platform in addressing metastatic melanoma and provide an in-depth immune profile study that validates its therapeutic potential. This work extends the knowledge from previous studies demonstrating DoriVac’s effectiveness in subcutaneous tumor models and highlights its capability against metastatic tumor models. This work highlights how the DoriVac platform could enhance the anti-tumor effects of FDA-approved immune checkpoint blockade antibodies in the clinic.

## Author contributions

This work was written by Anjali Rajwar*, Hawa Dembele*, Yang C. Zeng, and William M. Shih. Y.C.Z. and A.R. developed the idea for this manuscript. Y.C.Z., and A.R. planned experiments. A.R., Y.C.Z., H.D., A.R.G., A.V., M.S., and S.B. performed experiments. A.R., and H.D. drafted the manuscript. Y.C.Z., W.M.S. J.H.R and I.C.K. provided guidance on the project and edited the manuscript.

## Data and materials availability

The paper and supplementary materials contain the supporting data for this study.

## References

(1) Gershenwald, J. E.; Scolyer, R. A.; Hess, K. R.; Sondak, V. K.; Long, G. V.; Ross, M. I.; Lazar, A. J.; Faries, M. B.; Kirkwood, J. M.; McArthur, G. A.; Haydu, L. E.; Eggermont, A. M. M.; Flaherty, K. T.; Balch, C. M.; Thompson, J. F.; for members of the American Joint Committee on Cancer Melanoma Expert Panel and the International Melanoma Database and Discovery Platform. Melanoma Staging: Evidence-Based Changes in the American Joint Committee on Cancer Eighth Edition Cancer Staging Manual. CA. Cancer J. Clin. 2017, 67 (6), 472–492. 10.3322/caac.21409.

(2) Mishra, H.; Mishra, P. K.; Ekielski, A.; Jaggi, M.; Iqbal, Z.; Talegaonkar, S. Melanoma Treatment: From Conventional to Nanotechnology. J. Cancer Res. Clin. Oncol. 2018, 144 (12), 2283–2302. 10.1007/s00432-018-2726-1.

(3) Avula, L. R.; Grodzinski, P. Nanotechnology-Aided Advancement in the Combating of Cancer Metastasis. Cancer Metastasis Rev. 2022, 41 (2), 383–404. 10.1007/s10555-022-10025-7.

(4) Liu, J.; Fu, M.; Wang, M.; Wan, D.; Wei, Y.; Wei, X. Cancer Vaccines as Promising Immuno-Therapeutics: Platforms and Current Progress. J. Hematol. Oncol.J Hematol Oncol 2022, 15 (1), 28. 10.1186/s13045-022-01247-x.

(5) Jou, J.; Harrington, K. J.; Zocca, M.-B.; Ehrnrooth, E.; Cohen, E. E. W. The Changing Landscape of Therapeutic Cancer Vaccines—Novel Platforms and Neoantigen Identification. Clin. Cancer Res. 2021, 27 (3), 689–703. 10.1158/1078-0432.CCR-20-0245.

(6) Carvalho, T. Personalized Anti-Cancer Vaccine Combining mRNA and Immunotherapy Tested in Melanoma Trial. Nat. Med. 2023, 29 (10), 2379–2380. 10.1038/d41591-023-00072-0.

(7) Seeman, N. C. DNA in a Material World. Nature 2003, 421 (6921), 427–431. 10.1038/nature01406.

(8) Seeman, N. C. Nucleic Acid Junctions and Lattices. J. Theor. Biol. 1982, 99 (2), 237– 247. 10.1016/0022-5193(82)90002-9.

(9) Bachmann, M. F.; Jennings, G. T. Vaccine Delivery: A Matter of Size, Geometry, Kinetics and Molecular Patterns. Nat. Rev. Immunol. 2010, 10 (11), 787–796. 10.1038/nri2868.

(10) Pugh, G. C.; Burns, J. R.; Howorka, S. Comparing Proteins and Nucleic Acids for Next-Generation Biomolecular Engineering. Nat. Rev. Chem. 2018, 2 (7), 113–130. 10.1038/s41570-018-0015-9.

(11) Veneziano, R.; Moyer, T. J.; Stone, M. B.; Wamhoff, E.-C.; Read, B. J.; Mukherjee, S.; Shepherd, T. R.; Das, J.; Schief, W. R.; Irvine, D. J.; Bathe, M. Role of Nanoscale Antigen Organization on B-Cell Activation Probed Using DNA Origami. Nat. Nanotechnol. 2020, 15 (8), 716–723. 10.1038/s41565-020-0719-0.

(12) Hellmeier, J.; Platzer, R.; Eklund, A. S.; Schlichthaerle, T.; Karner, A.; Motsch, V.; Schneider, M. C.; Kurz, E.; Bamieh, V.; Brameshuber, M.; Preiner, J.; Jungmann, R.; Stockinger, H.; Schütz, G. J.; Huppa, J. B.; Sevcsik, E. DNA Origami Demonstrate the Unique Stimulatory Power of Single pMHCs as T Cell Antigens. Proc. Natl. Acad. Sci. 2021, 118 (4), e2016857118. 10.1073/pnas.2016857118.

(13) Zeng, Y. C.; Young, O. J.; Wintersinger, C. M.; Anastassacos, F. M.; MacDonald, J. I.; Isinelli, G.; Dellacherie, M. O.; Sobral, M.; Bai, H.; Graveline, A. R.; Vernet, A.; Sanchez, M.; Mulligan, K.; Choi, Y.; Ferrante, T. C.; Keskin, D. B.; Fell, G. G.; Neuberg, D.; Wu, C. J.; Mooney, D. J.; Kwon, I. C.; Ryu, J. H.; Shih, W. M. Fine Tuning of CpG Spatial Distribution with DNA Origami for Improved Cancer Vaccination. Nat. Nanotechnol. 2024, 1–11. 10.1038/s41565-024-01615-3.

(14) Zeng, Y. C.; Young, O. J.; Si, L.; Ku, M. W.; Isinelli, G.; Rajwar, A.; Jiang, A.; Wintersinger, C. M.; Graveline, A. R.; Vernet, A.; Sanchez, M.; Ryu, J. H.; Kwon, I. C.; Goyal, G.; Ingber, D. E.; Shih, W. M. DNA Origami Vaccine (DoriVac) Nanoparticles Improve Both Humoral and Cellular Immune Responses to Infectious Diseases. bioRxiv January 2, 2024, p 2023.12.29.573647. 10.1101/2023.12.29.573647.

(15) Shen, F.; Wang, H.; Liu, Z.; Sun, L. DNA Nanostructures: Self-Adjuvant Carriers for Highly Efficient Subunit Vaccines. Angew. Chem. 2024, 136 (2), e202312624. 10.1002/ange.202312624.

(16) Qu, Y.; Shen, F.; Zhang, Z.; Wang, Q.; Huang, H.; Xu, Y.; Li, Q.; Zhu, X.; Sun, L. Applications of Functional DNA Materials in Immunomodulatory Therapy. ACS Appl. Mater. Interfaces 2022, 14 (40), 45079–45095. 10.1021/acsami.2c13768.

(17) Oktay, E.; Alem, F.; Hernandez, K.; Girgis, M.; Green, C.; Mathur, D.; Medintz, I. L.; Narayanan, A.; Veneziano, R. DNA Origami Presenting the Receptor Binding Domain of SARS-CoV-2 Elicit Robust Protective Immune Response. Commun. Biol. 2023, 6 (1), 1–11. 10.1038/s42003-023-04689-2.

(18) Liu, S.; Jiang, Q.; Zhao, X.; Zhao, R.; Wang, Y.; Wang, Y.; Liu, J.; Shang, Y.; Zhao, S.; Wu, T.; Zhang, Y.; Nie, G.; Ding, B. A DNA Nanodevice-Based Vaccine for Cancer Immunotherapy. Nat. Mater. 2021, 20 (3), 421–430. 10.1038/s41563-020-0793-6.

(19) Choi, Y.; Cho, B. K.; Seok, S. H.; Kim, C.; Ryu, J. H.; Kwon, I. C. Controlled Spatial Characteristics of Ligands on Nanoparticles: Determinant of Cellular Functions. J. Controlled Release 2023, 360, 672–686. 10.1016/j.jconrel.2023.07.020.

(20) Zeng, Y. C.; Young, O. J.; Wintersinger, C. M.; Anastassacos, F. M.; MacDonald, J. I.; Isinelli, G.; Dellacherie, M. O.; Sobral, M.; Bai, H.; Graveline, A. R.; Vernet, A.; Sanchez, M.; Mulligan, K.; Choi, Y.; Ferrante, T.C.; Keskin, D.B.; Fell, G.G.; Neuberg, D.; Wu, C.J.; Mooney, D.J.; Kwon, I.C.; Ryu, J.H.; Shih, W.M. Optimizing CpG Spatial Distribution with DNA Origami for Th1-Polarized Therapeutic Vaccination. bioRxiv August 27, 2022, p 2022.06.08.495340. 10.1101/2022.06.08.495340.

(21) Wintersinger, C. M.; Minev, D.; Ershova, A.; Sasaki, H. M.; Gowri, G.; Berengut, J. F.; Corea-Dilbert, F. E.; Yin, P.; Shih, W. M. Multi-Micron Crisscross Structures Grown from DNA-Origami Slats. Nat. Nanotechnol. 2023, 18 (3), 281–289. 10.1038/s41565-022-01283-1.

(22) Buchberger, A.; Simmons, C. R.; Fahmi, N. E.; Freeman, R.; Stephanopoulos, N. Hierarchical Assembly of Nucleic Acid/Coiled-Coil Peptide Nanostructures. J. Am. Chem. Soc. 2020, 142 (3), 1406–1416. 10.1021/jacs.9b11158.

(23) Dudek, A. M.; Martin, S.; Garg, A. D.; Agostinis, P. Immature, Semi-Mature, and Fully Mature Dendritic Cells: Toward a DC-Cancer Cells Interface That Augments Anticancer Immunity. Front. Immunol. 2013, 4. 10.3389/fimmu.2013.00438.

(24) Reis e Sousa, C. Dendritic Cells in a Mature Age. Nat. Rev. Immunol. 2006, 6 (6), 476– 483. 10.1038/nri1845.

(25) Ma, D. Y.; Clark, E. A. The Role of CD40 and CD154/CD40L in Dendritic Cells. Semin. Immunol. 2009, 21 (5), 265–272. 10.1016/j.smim.2009.05.010.

(26) Oh, S. A.; Wu, D.-C.; Cheung, J.; Navarro, A.; Xiong, H.; Cubas, R.; Totpal, K.; Chiu, H.; Wu, Y.; Comps-Agrar, L.; Leader, A. M.; Merad, M.; Roose-Germa, M.; Warming, S.; Yan, M.; Kim, J. M.; Rutz, S.; Mellman, I. PD-L1 Expression by Dendritic Cells Is a Key Regulator of T-Cell Immunity in Cancer. Nat. Cancer 2020, 1 (7), 681–691. 10.1038/s43018-020-0075-x.

(27) Peng, Q.; Qiu, X.; Zhang, Z.; Zhang, S.; Zhang, Y.; Liang, Y.; Guo, J.; Peng, H.; Chen, M.; Fu, Y.-X.; Tang, H. PD-L1 on Dendritic Cells Attenuates T Cell Activation and Regulates Response to Immune Checkpoint Blockade. Nat. Commun. 2020, 11 (1), 4835. 10.1038/s41467-020-18570-x.

(28) Tang, H.; Wang, Y.; Chlewicki, L. K.; Zhang, Y.; Guo, J.; Liang, W.; Wang, J.; Wang, X.; Fu, Y.-X. Facilitating T Cell Infiltration in Tumor Microenvironment Overcomes Resistance to PD-L1 Blockade. Cancer Cell 2016, 29 (3), 285–296. 10.1016/j.ccell.2016.02.004.

(29) Gordon, S.; Taylor, P. R. Monocyte and Macrophage Heterogeneity. Nat. Rev. Immunol. 2005, 5 (12), 953–964. 10.1038/nri1733.

(30) Hong, S. T.; You, D. G.; Jo, M.; Kim, C. H.; Choi, Y.; Kim, C.; Park, J. H.; Kim, K.; Kwon, I. C.; Ryu, J. H. Imaging of Tumor-Associated Macrophages Using near-Infrared Fluorophore-Conjugated Dextran-Sulfate Nanoparticles. Macromol. Res. 2023, 31 (12), 1113– 1124. 10.1007/s13233-023-00201-1.

(31) Perez, C.; Botta, C.; Zabaleta, A.; Puig, N.; Cedena, M.-T.; Goicoechea, I.; Alameda, D.; San José-Eneriz, E.; Merino, J.; Rodríguez-Otero, P.; Maia, C.; Alignani, D.; Maiso, P.; Manrique, I.; Lara-Astiaso, D.; Vilas-Zornoza, A.; Sarvide, S.; Riillo, C.; Rossi, M.; Rosiñol, L.; Oriol, A.; Blanchard, M.-J.; Rios, R.; Sureda, A.; Martin, J.; Martinez, R.; Bargay, J.; de la Rubia, J.; Hernandez, M.-T.; Martinez-Lopez, J.; Orfao, A.; Agirre, X.; Prosper, F.; Mateos, M.-V.; Lahuerta, J.-J.; Blade, J.; San-Miguel, J. F.; Paiva, B.; on behalf of the Grupo Español de Mieloma/Programa para el Estudio de la Terapéutica en Hemopatías Malignas Cooperative Study Group. Immunogenomic Identification and Characterization of Granulocytic Myeloid-Derived Suppressor Cells in Multiple Myeloma. Blood 2020, 136 (2), 199–209. 10.1182/blood.2019004537.

(32) Vignali, D. A. A.; Collison, L. W.; Workman, C. J. How Regulatory T Cells Work. Nat. Rev. Immunol. 2008, 8 (7), 523–532. 10.1038/nri2343.

(33) Aktas, E.; Kucuksezer, U. C.; Bilgic, S.; Erten, G.; Deniz, G. Relationship between CD107a Expression and Cytotoxic Activity. Cell. Immunol. 2009, 254 (2), 149–154. 10.1016/j.cellimm.2008.08.007.

