## Supplementary Information for "Validation of DoriVac (DNA origami vaccine) efficacy in a metastatic melanoma model"

| Antibodies | Fluorophores | Biolegend Catalog No. |  | Antibodies | Fluorophores | Biolegend Catalog No. |
| --- | --- | --- | --- | --- | --- | --- |
| CD11c | BV570 | 117331 |  | CD3 | BV585 | 100225 |
| CD40 | PE | 124610 |  | CD4 | PE | 100408 |
| CD86 | PE/CY7 | 105014 |  | CD8 | APC/CY7 | 100714 |
| MHC-II | BV421 | 107632 |  | CD25 | AF700 | 102024 |
| PD-L1 | BV711 | 124319 |  | CD44 | AF700 | 103026 |
| DEC205 | PerCP/Cy5.5 | 138208 |  | FoxP3 | BV421 | 126419 |
| Viability | Zombie UV | 423108 |  | IFNγ | BV510 | 505841 |
| CD11b | BV605 | 101257 |  |  |  |  |
| Gr-1 | FITC | 108406 |  |  |  |  |
| F4/80 | PE594 | 123146 |  |  |  |  |
| CD206 | BV650 | 141723 |  |  |  |  |
| NK1.1 | PE | 156504 |  |  |  |  |
| CD64 | AF488 | 305010 |  |  |  |  |
| T-bet | BV421 | 644832 |  |  |  |  |
| TNFα | PE594 | 506346 |  |  |  |  |
| CD107a | AF488 | 121608 |  |  |  |  |

**Table S1.** Antibodies used to stain the lymphocytes for flow cytometry collection.

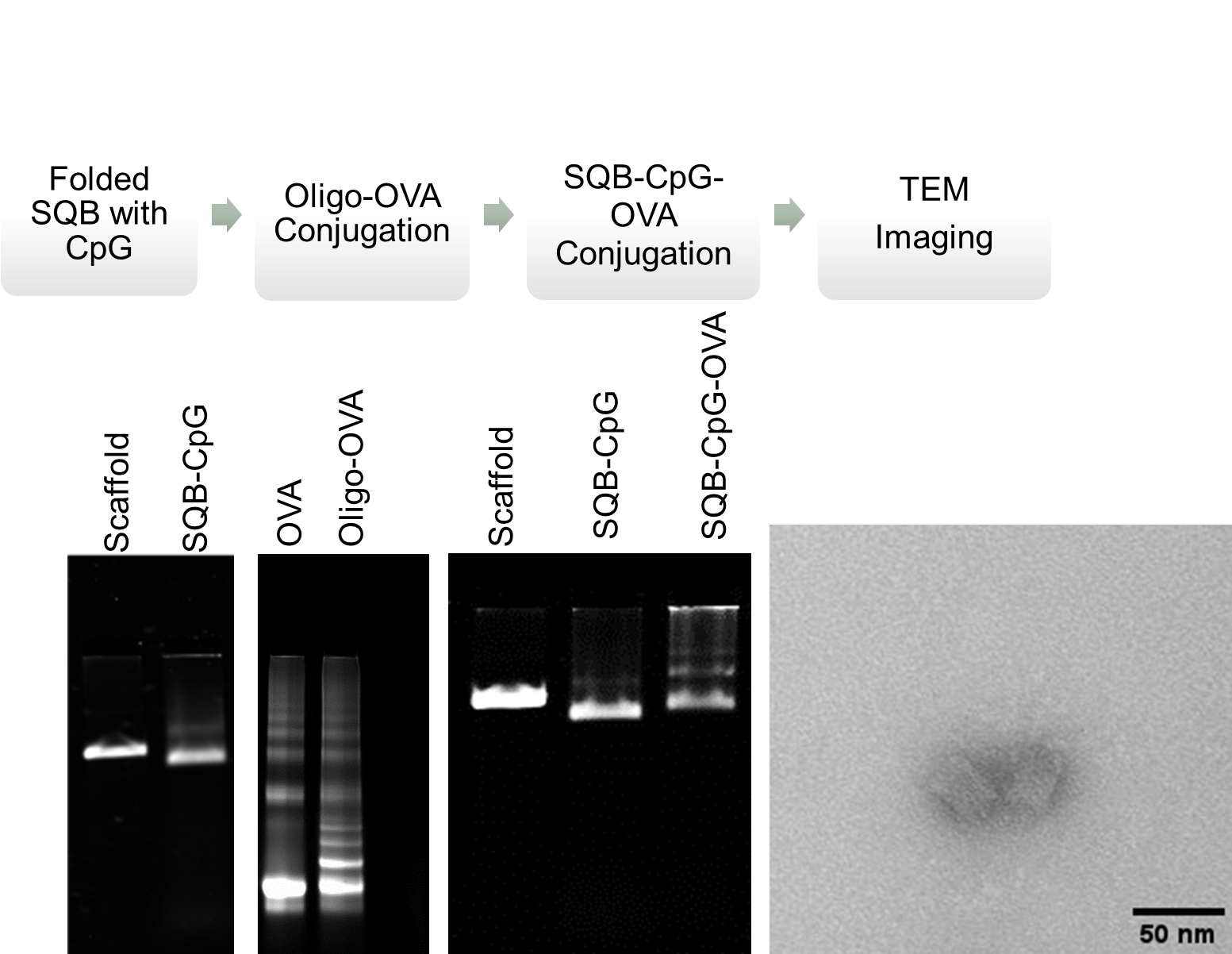

Figure 1: Fabrication of SQB DoriVac for B16OVA metastasis study. SQB nanoparticles were assembled with CpG oligonucleotides using the p8634 scaffold. An SDS-PAGE gel demonstrates the conjugation of the OVA antigen with the anti-handle oligonucleotide using an SMCC crosslinker. An agarose gel shows the conjugation of the OVA antigen with SQB nanoparticles through handle-anti-handle conjugation chemistry. Representative transmission electron microscopy image of SQB DoriVac.

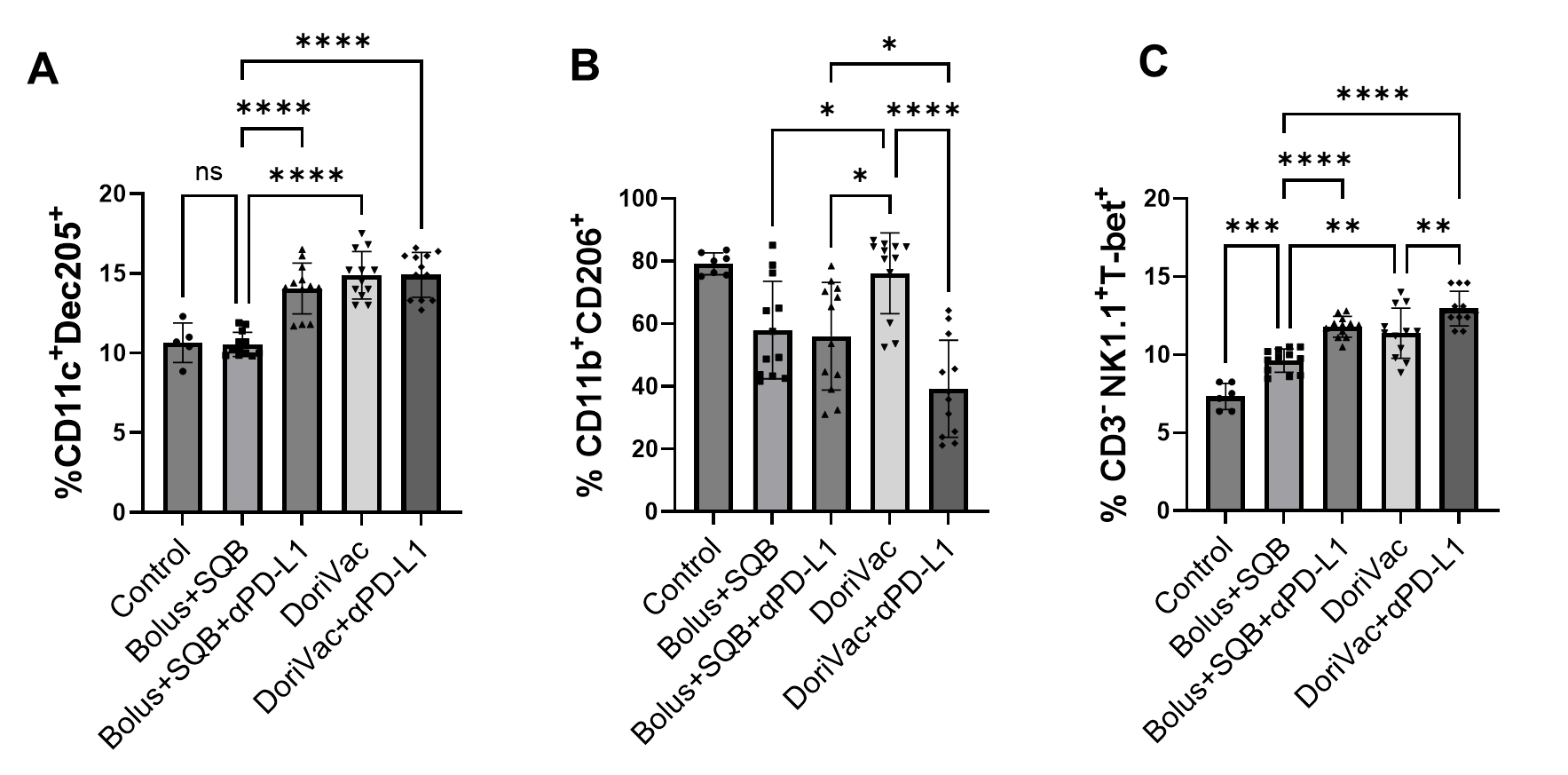

Figure 2: Effect of DoriVac treatment on innate immune cells from the lymph nodes in B16OVA model. (A) Quantification of CD11c^+^Dec205^+^ subset of dendritic cells that co-express CD11c and Dec205. (B) Quantification of CD11b^+^CD206^+^ that are indicative of M2 macrophages. (C) Quantification of CD3^-^NK1.1^+^Tbet^+^ activated NK cells expressing T-bet transcription factor. Data was collected from six mice within each group. The control groups received no treatment. Error bars indicate the mean with the associated standard deviation. Statistical analysis was performed using one-way ANOVA followed by Tukey’s post hoc multiple comparison test. ****p < 0.0001, **p < 0.01, *p < 0.05, ns: non-significant. Samples from each mouse were duplicated for flow cytometry, except for the control group. Therefore, the control group has n=6 while the other groups each have n=12.

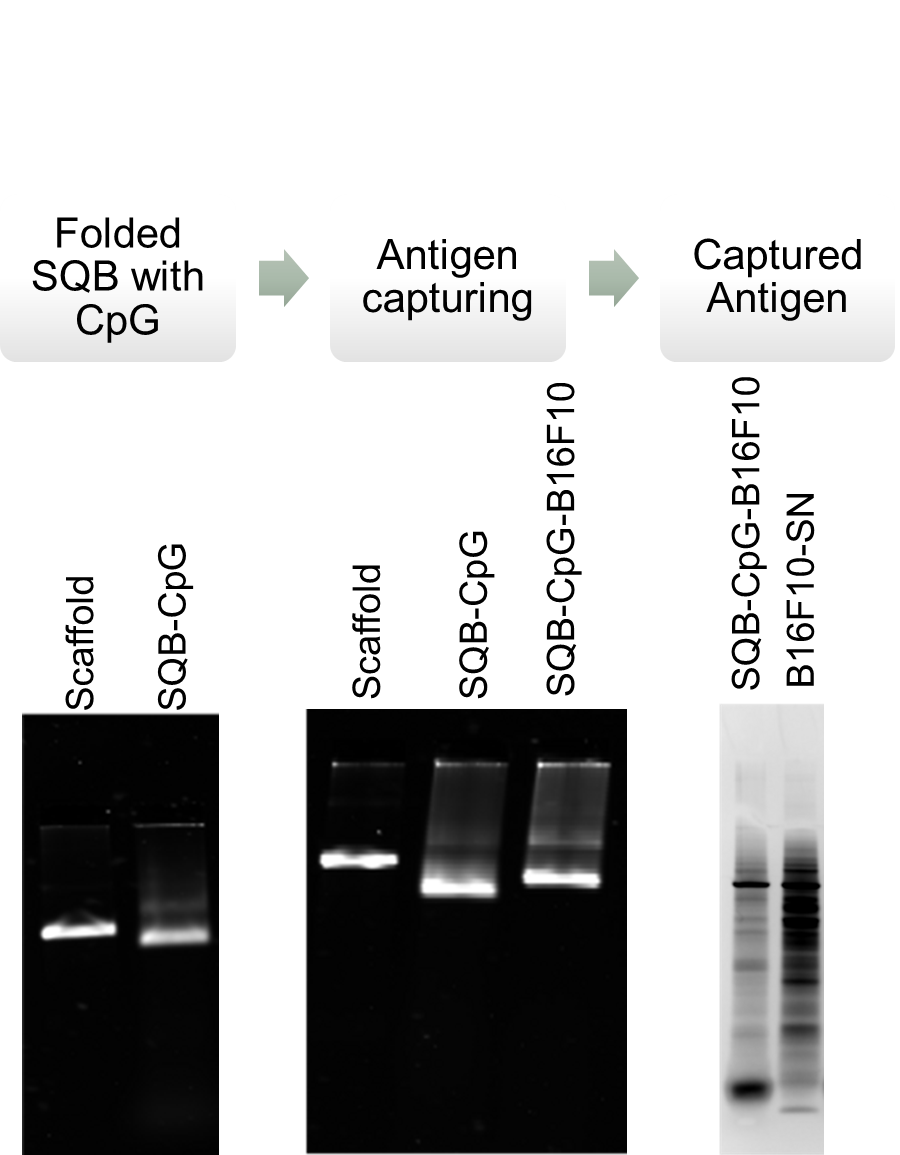

Figure 3: Fabrication of SQB DoriVac for B16F10 study. SQB nanoparticles were assembled with CpG oligonucleotides using the p8634 scaffold. These SQB nanoparticles were then co-cultured with B16F10 tumor cell supernatant to form SQB DoriVac by capturing antigens. The quantity of captured antigen was confirmed through DNase digestion, which degraded the SQB origami particles and left the captured antigens, which were subsequently stained using silver staining. SN: supernatant.

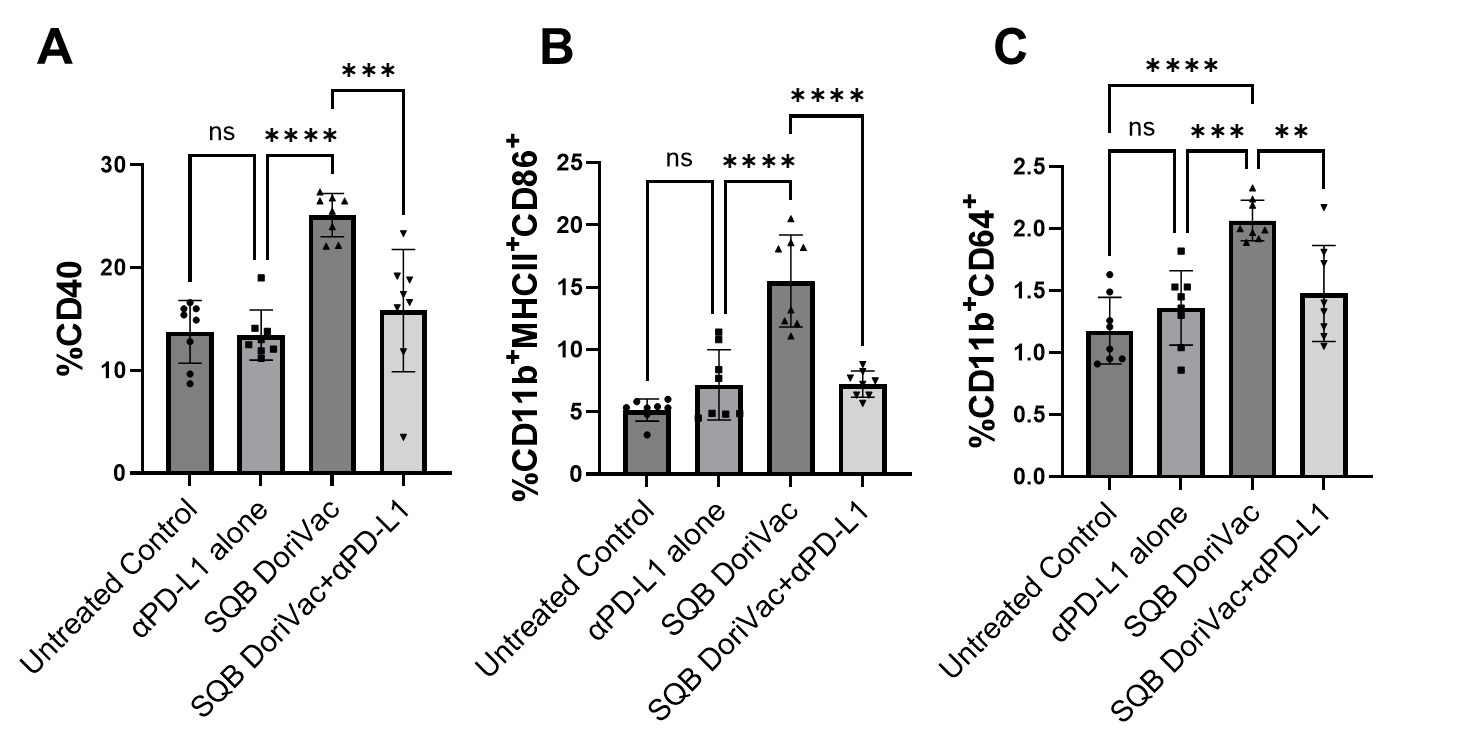

Figure 4: DoriVac treatment activates DC and Macrophage Populations from lymph nodes in the B16F10 Model (A) Quantification of CD40 cells with αPD-L1 and DoriVac monotherapy and their combination. (B) Quantification of CD11b^+^MHCII^+^CD86^+^ dendritic cells (C) Quantification of CD11b^+^CD64^+^ macrophages. Data was collected from four mice within each group. The control groups received no treatment. Error bars indicate the mean with the associated standard deviation. Statistical analysis was performed using one-way ANOVA followed by Tukey’s post hoc multiple comparison test. (****p < 0.0001, ***p < 0.001, **p < 0.01, ns: non-significant. Samples from each mouse were duplicated for flow cytometry therefore each group has n=8.

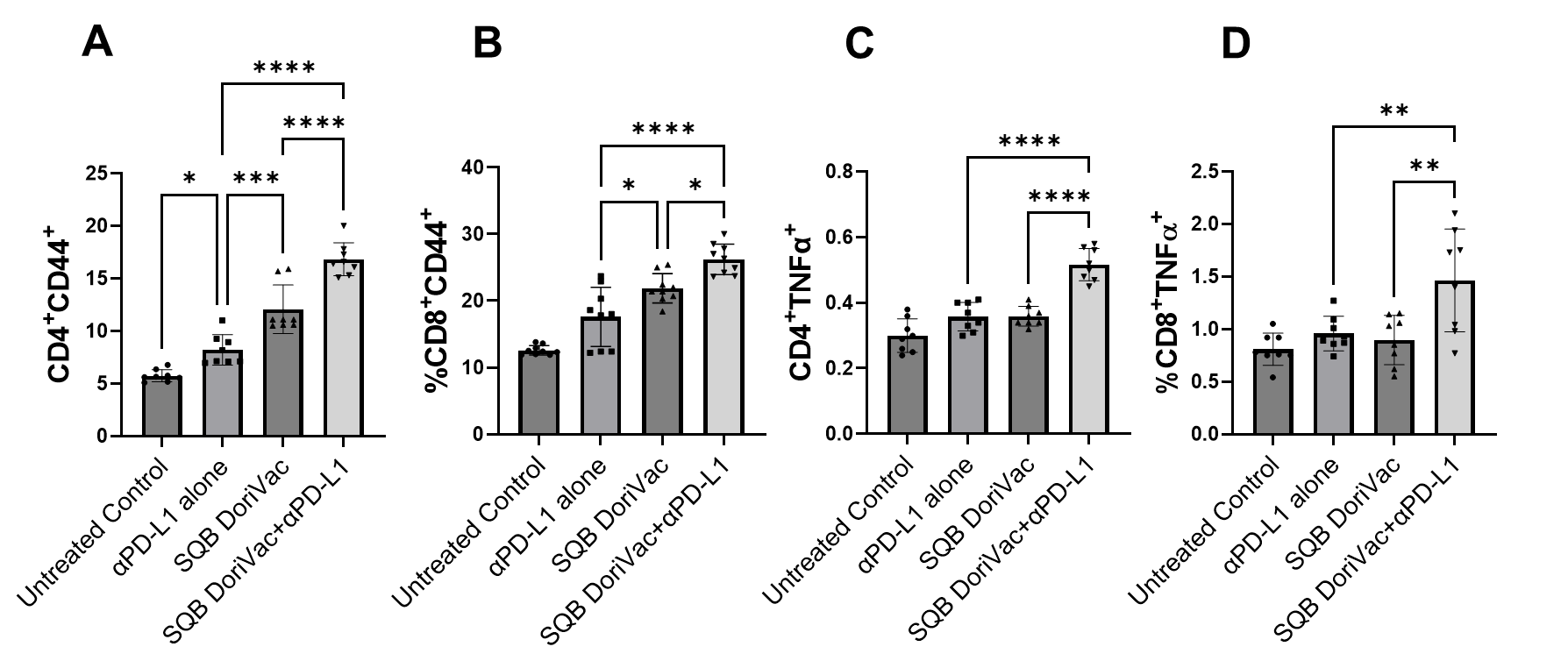

Figure 5: DoriVac in combination with αPD-L1 activates CD4 and CD8 T cells. (A) The percentages of CD4^+^CD44^+^ T cells represent a subset of memory helper CD4 T cells. (B) Percentage of memory cytotoxic CD8 T cells. (C) Percentages of CD4^+^ helper T-cells producing TNFα^+^ from mouse lymph nodes. (D) Percentages of CD8^+^ T-cells producing TNFα^+^ from mouse lymph nodes. Data was collected from four mice in each group. The untreated control groups received no treatment. Error bars indicate the mean with the associated standard deviation. Statistical analysis was performed using one-way ANOVA followed by Tukey’s post hoc multiple comparison test. (****p < 0.0001, ***p < 0.001, **p <0.01, *p<0.05, ns: non-significant. Samples from each mouse were duplicated for flow cytometry therefore each group has n=8.
